# autoFISH - a modular toolbox for sequential smFISH experiments

**DOI:** 10.1101/2024.11.14.623566

**Authors:** Christian Weber, Thomas Defard, Mickael Lelek, Hugo Laporte, Ayan Mallick, José-Arturo Londoño-Vallejo, Thomas Walter, Charles Fouillade, Maria Isabella Gariboldi, Florian Mueller

**Author notes:** These authors contributed equally to this work (Christian Weber, Thomas Defard, Florian Mueller).

## Abstract

Fluorescence in situ hybridization (FISH) allows for spatial and quantitative profiling of gene expression by visualizing individual RNA molecules. Here, we introduce automated FISH (autoFISH), a comprehensive toolbox to conduct automated single molecule FISH (smFISH) experiments that is both cost-effective and versatile. This includes detailed plans for constructing the necessary equipment, open-source software for control, reliable experimental protocols, and analysis workflows based on our FISH-quant analysis package. Validation experiments with both cell lines and tissue samples confirmed the system’s robustness. We demonstrate standard and amplified smFISH, along with a modified protocol for tissue clearing that enhances nuclear retention while preserving background reduction efficiency.

## Introduction

Transcription is a fundamental process in biology and plays a pivotal role in development and disease. Individual cells display considerable heterogeneity in their transcriptional profiles, both in RNA abundance and intracellular RNA localization. Accurately quantifying the number of RNAs within cells and their locations provides a precise readout of a cell’s molecular state and enhances our understanding of RNA regulation. Obtaining such measurements requires spatially resolved RNA detection in the native cellular environment with sub-cellular accuracy.

Here, single-molecule FISH (smFISH) is widely regarded as the gold standard due to its ability to detect individual RNA molecules with high sensitivity and its precise localization within the sub-micrometer range (Pichon et al., 2018). Various adaptations of this method have been developed, all based on the principle of targeting RNAs with multiple oligonucleotide fluorescent probes, either directly labeled or indirectly labeled via secondary probes. RNAs then appear as diffraction-limited spots in fluorescence microscopy images and their positions can be measured with high spatial accuracy using dedicated analysis workflows (Defard et al., 2024a). When combined with cell segmentation, the RNA abundance in each cell can be estimated, and statistical analysis can be used to quantitatively describe subcellular RNA localization (Chouaib et al., 2020). Importantly, smFISH offers insights into the transcriptional states of cells while preserving their spatial context, enabling large-scale measurements such as cell-to-cell communication, cell composition, and extensive tissue gradients (Palla et al., 2022).

A significant limitation of the original smFISH methods is that the number of spectrally distinguishable fluorophores limits the number of different RNA species that can be simultaneously visualized. To overcome this limitation, new methods have been developed that rely on iterative hybridizations, i.e. alternating cycles of probe hybridization, imaging, and signal removal (Pichon et al., 2018). A general feature of these approaches is that the primary probes carry one or several readout sequences that can be revealed with a secondary complementary, readout oligo. Usually, primary probes against all RNA targets are hybridized on the bench, while the readout oligos are hybridized directly on the microscope with an automated fluidic system. Signal removal can be accomplished through various methods, the most commonly used are: stripping with high stringency buffers (Lohoff et al., 2022), fluorophore removal by chemical cleavage (Moffitt et al., 2016b), or displacement with a tertiary stripping oligo (Mateo et al., 2021). In sequential smFISH experiments, the number of targeted species is in linear correlation with the number of iterations and fluorophores used. A dramatic increase in the number of RNA species throughput can be achieved by multiplexed barcoding (Pichon et al., 2018). Here, each RNA species carries multiple, distinct readout sequences allowing its detection in different hybridization rounds, defining a unique detection barcode. Importantly, some readout sequences are shared by different RNA species. In each hybridization round, RNAs from different genes are thus visualized together but never the same. Molecular identity of each detected RNA is then determined in a post-processing step by decoding these barcodes. This combinatorial approach combined with error correction strategies permits the detection of hundreds to thousands of different RNA species (Eng et al., 2019; Zhang et al., 2021). While several commercial systems offer convenient turnkey solutions, they are costly, both in terms of instrumentation and reagents, and lack flexibility. Exploratory analyses, time-courses, dose-response studies, and replicates thus demand substantial financial resources. Additionally, their ease of use comes at the expense of flexibility, both in microscopy (e.g., imaging modality) and experimental protocols (e.g., adding signal amplification steps). While large RNA panels are often necessary, not all studies require such depth; smaller, more flexible panels can sometimes be sufficient, e.g. a small probe panel with 10 genes can be sufficient to resolve main cell types in tumor cryosections (Rademacher et al., 2024). Therefore, we believe that home-built-solutions, both in instrumentation and experimental workflows, remain an important option for many research questions.

To address this need, we propose autoFISH, an end-to-end open-source workflow comprising hardware, software and experimental protocols for automated smFISH experiments designed for flexibility and affordability. The autoFISH toolbox relies on a versatile and affordable fluidic system, for which we provide detailed building plans. Flexibility comes in several flavors. First, the imaging modality is not fixed since the fluidics system required for sequential hybridizations can be mounted on different kinds of microscopes depending on the desired application. Second, protocols can be easily modified depending on the experimental need, e.g. for inclusion of signal amplification or tissue clearing. Lastly, since probe design and synthesis can be controlled, smaller scale experiments can easily be performed.

We demonstrate the applicability of autoFISH on two experimental datasets. Firstly, HeLa cells, which we use to benchmark the performance of the protocol and illustrate its flexibility by alternating between standard and amplified FISH experiments with SABER (Kishi et al., 2019), and secondly, mouse lung tissue, which we use to visualize several marker genes from our earlier study (Curras-Alonso et al., 2023). Here, we introduce a modified protocol for tissue clearing with an improved nuclear signal. In summary, we provide a complete toolbox to establish automated, sequential smFISH experiments at a lower price tag.

## Methods

In this section we provide an overview of the methods. For detailed, up-to-date information, readers are referred to the following dedicated GitHub repositories: https://github.com/fish-quant/autofish (for the fluidics system and experimental protocols) and https://github.com/fish-quant/autofish-analysis (analysis code).

### Instrumentation and control

AutoFISH relies on an automated fluidics system for buffer exchanges. A Python control software was developed to control the fluidics system and communicate with the microscope to launch acquisitions. AutoFISH is implemented as a light-weight Python package with only minimal dependencies. It uses PySerial for communication with various hardware components, and its modular design allows for the easy addition of new components, like pumps or valves from different suppliers. AutoFISH can be controlled with a graphical user interface.

### Microscopy

AutoFISH does not provide the possibility to directly control different microscopes, but provides different options to trigger acquisitions. This renders autoFISH extremely flexible as it is usable on a range of microscopes with minimal adaptation. At the time of publication, the following options are available, but new ones can be easily added:

Pycromanager (Pinkard et al., 2021) - suitable for microscopes that are controlled by micromanger. Here, the user can define a settings file with the relevant acquisition parameters, as well as a position list. autoFISH will then create a multi-D acquisition event that permits acquiring images after each hybridization round.

Synchronization with a text file; either the file content, e.g a switch from 0 to 1, or the existence of a file can be used to start an acquisition. Once the acquisition is completed the file content can be set back to 0, or the file can be deleted, signaling to autoFISH that the next hybridization round can be started.

Communication with TTL trigger: suitable for microscopes that can integrate incoming and outgoing TTL trigger events in their acquisition routines. In autoFISH, we use an Arduino to send/receive such TTL triggers: a TTL trigger is sent to start an acquisition, and incoming TTL trigger is used to indicate that the acquisition is terminated and that the next hybridization rounds can be started.

### Fluidics system

We use a custom-built fluidics system combining elements from published approaches (merFISH and ORCA (Bintu et al., 2018; Moffitt and Zhuang, 2016)). Detailed building plans and part lists can be found on GitHub (https://github.com/fish-quant/autofish). The system consists of the following main components, which are widely available and affordable:

Several elements guarantee reliable flow of different buffers: a peristaltic pump for accurate flow rates, a bubble trap connected to a vacuum pump to remove air bubbles, and a flow sensor to monitor flow and detect any potential issues such as obstructed fluidic lines or large air bubbles.

Buffers are stored either in deep-well 96-well plates (each well holding up to 2 mL of buffer) or in syringes. The former usually contains hybridization buffers that are used only once and the syringes general-purpose buffers for washing and imaging. The plates are placed on a pipette robot built from a CNC milling machine. These robots can be easily adapted by including a 3D-printed insert in the spindle holder to accommodate a needle connected to a fluidic line. The needle can aspirate liquid from the multi-well plates. A computer-controlled valve selects buffers from different inlets, either from the fluidic line connected to the pipette robot or the syringes.

Lastly, several sample support systems are presented. The choice depends on the number of samples and sample characteristics. First, temperature controlled imaging chamber Bioptechs FCS2 for individual samples or tissue samples Bioptechs FCS2 chamber (exist for inverted and upright microscopes). Second, Ibidi glass bottom, 6-chanel microslide to run multiple samples in parallel. To ensure temperature control, these channel slides are placed in Okolab Stage Top chambers. These channel slides can be easily integrated in the fluidic system having female Luer adaptor inlets. To guarantee that each channel is appropriately perfused with a buffer, a distribution valve is used so that each channel can be flushed separately. Note that the Hamiliton MVP valves that were used can be used in either flow direction.

The total price for the complete fluidics system is less than 5,000 EUR excluding sample support. The FCS2 imaging chamber costs around 5,000 EUR, whereas the disposable Ibidi slides can be readily used for cultured cell lines and cost on the order of 15 euros each.

### Sequential smFISH

We provide detailed protocols here https://github.com/fish-quant/autofish.

#### General points

In experiments performed on the fluidic system, we used the less toxic Ethylene Carbonate (EC) instead of formamide, as previously described (Moffitt et al., 2016a). In most previous studies, hybridizations on the fluidics system were done without Dextran. For some experiments, e.g. signal amplification with SABER, we found that using even low dextran helps to reduce oligo concentrations, improve signal and reduce background. With our peristaltic pump (Ismatec Reglo Digital), we can use up to 5% dextran without detrimental impact on flow rates. Note, it is important to filter all buffers (0.22µm filters) to avoid clogging the fluidic lines. The imaging buffer contains enzymes. To maintain a low temperature throughout the experiment, an isothermal chamber with a cold block is used. In the syringe, a layer of mineral oil on top of the imaging buffer prevents oxygen from diffusing into the buffer.

#### Primary oligos

Oligos against the target genes can be designed with a range of different tools (Defard et al., 2024a). Here, we used Oligostan (Tsanov et al., 2016), a computational tool we built in-house for bioinformatic design of RNA probes. The oligo synthesis method depends on the number of required oligos. For small scale studies involving only a few hundred oligos, we use oligo pools provided by Integrated DNA technologies (IDT) at a concentration of 50 pmol/oligo. We found that these pools can be used without amplification, and are sufficient for several hundred hybridization rounds when hybridizing 12mm coverslips. Individual pools can be ordered for each gene, which then also allows custom assembly of hybridization panels for specific questions. For larger oligo numbers, oPools that require amplification before usage are used (Moffitt and Zhuang, 2018).

#### smFISH readout and stripping oligos

In this study, we employed the strategy outlined by (Bintu et al., 2018), which utilizes three different oligos: primary oligos, secondary imager-conjugated oligos, and displacement oligos. For each gene, 20-50 primary oligos are designed (see above) and consist of two segments. The first segment is a DNA stretch complementary to a portion of the target gene’s RNA. The second segment is a ‘readout’ overhang sequence. This readout is shared by all oligos targeting the same RNA. Secondary oligos have regions complementary to the primary oligo readout region. They also include a sequence pre-hybridized to one or multiple fluorophore-conjugated imager probes, which can be different colors depending on the assigned fluorophore. A third region, the so-called toehold sequence, is unbound. To remove the signal, the third displacement oligo can be used. This oligos peels off the secondary oligo and the resulting oligo duplex can be washed out. In the actual experiment, all primary oligos can be hybridized simultaneously during a primary incubation step, while secondary oligos are hybridized sequentially to image each RNA species consecutively. Multiple RNA species can be imaged simultaneously using secondary probes conjugated to spectrally distinct fluorophores. Displacement probes can be used to remove readout probes from previous rounds, facilitating this multiplexing approach and enabling experiments to be conducted with low recurrent costs.

We observed that some provided readout sequences yielded lower signals when used for RNA smFISH and developed a simple strategy to systematically test their quality. In short, we used a probe-set against a housekeeping gene carrying a validated readout sequence (RO1). We then design an intermediate oligo with a 5’ region that has a reverse sequence complementary to RO1 and a 3’ region with a reverse sequence complementary to the readout to be tested. The to be tested readout can then be hybridized with an imager oligo and visualized. This approach eliminates the need to synthesize a reference oPool for each new ReadOut, requiring only a simple oligo instead. For an up-to-date list of validated readout and imager sequences, please refer to the resources provided at https://github.com/fish-quant/autofish.

#### Saber

SABER provides a simple way to amplify the signal in a programmable way (Kishi et al., 2019). For us, amplification up to 10X works well without large aggregations. By coupling the amplification hairpins and the ORCA readout sequences, specific hybridization rounds can be amplified, and also stripped. When optimizing SABER, we found that due to the signal amplification often abundant non-specific dots can be seen that stem from either non-specific primary probes or amplicons. To reduce this background, we found that a 3-step hybridization works best to reduce this background by optimizing strigency for each step Firstly, primary probes are hybridized on the bench and we use a more stringent buffer (1X SSC, 30% formamide, 10% dextran) instead (2X SSC, 20% formamide). Then, the amplicons are hybridized on the fluidics system for each round with 1xSSC, 20% EC, 5% dextran. Lastly, the imaging oligos are hybridized on the fluidics system (2X SSC, 10% EC, 5% dextran). The TM of these oligos is relatively low since they contain no G, and a more stringent buffer leads to loss in signal intensity. The used low concentration of dextran does not impact the buffer exchange, but allows to reduce the concentration of the imaging oligos.

#### Clearing

Sample clearing reduces background signal in smFISH by removing tissue autofluorescence and reducing off-target binding of probes to lipids and proteins (Moffitt et al., 2016a). Briefly, RNAs are first anchored to a polyacrylamide matrix using poly-A targeting oligos, or epoxide (GMA). Once the matrix has solidified, tissues are incubated at 37°C overnight in a Proteinase K solution (20mg/ml). The next day, tissues are washed, and hybridization of readout oligos is performed.

### Validation experiments

#### Sequential smFISH in HeLa cells

To validate the autoFISH procedure, HeLa cells were first cultured in DMEM supplemented with 10% FBS and 1% penicillin streptomycin. The cells were then fixed with a 4% PFA solution and permeabilized with 70% ethanol at -20°C overnight. After rehydration, the samples were hybridized with primary probes against XPO1 and KIF1C overnight. Following mounting in the appropriate support, the samples were installed on the fluidics system. A series of 20 alternating hybridizations and stripping of XPO1 and KIF1C was then conducted on the fluidic system. This allowed for the monitoring of signal quality over time, observation of probe dehybridization, and tracking of the signal quality throughout the experiment.

#### Mouse lung tissue

To validate the hybridization of several genes on a section of fixed and cleared tissue, autoFISH was performed on mouse lung tissue. Briefly, mouse lungs were inflated with cold 4% paraformaldehyde (PFA), fixed overnight at 4°C, cryopreserved in 30% sucrose and embedded in optimal cutting temperature (OCT) compound before sectioning (Curras-Alonso et al., 2023).

### Analysis

After completing the experiment, the first step in the analysis is to segment the acquired images into .tiff files, each containing the entire Z-stack for a field of view and a hybridization round. During the experiment, alignment between each field of view may drift between rounds. To correct this, we use the SimpleITK package (Beare et al., 2018), which employs similarity metrics between images across rounds to achieve alignment.

For analysis of cell-culture experiments, spots are detected with Big-FISH (Imbert et al., 2022). For lung tissue experiments, spot detection was performed with the recent deep learning approach U-FISH (Xu et al., 2024). Spot quality metrics, such as signal-to-noise ratio, background, and spot intensity, were computed using Big-FISH (Imbert et al., 2022). Nuclei segmentation on DAPI and cell segmentation, when cell staining was present, are performed with Cellpose (Pachitariu and Stringer, 2022). Finally, all fields of view are stitched using the stitching ImageJ plugin (Preibisch et al., 2009). The final step is to generate the spatially resolved gene-by-cell expression matrix. However, since cell boundary staining is not always available in our experiments, the cell expression matrix is calculated with our recently published point cloud segmentation method, ComSeg (Defard et al., 2024b). To perform cell type calling, we calculated the cosine distance between the cell expression vectors and the median centroids of each cell type, as defined in the reference scRNA-seq data from (Curras-Alonso et al., 2023). These gene markers allow to classify five distinct cell types: Alveolar Macrophage (AM) with *Chil3*, Alveolar Type 1 (AT1) with *Rtkn2*, Alveolar Type 2 (AT2) with *Lamp3*, and Epithelial Cells with *Pecam1 and Ptprb*. Cells are assigned to their nearest cell type cluster based on cosine distance. Given the limitations of our gene markers, not every cell can be classified. Therefore, we only classify cells with a cosine distance to the nearest cell type centroid below 0.8 and more than 10 detected RNAs.

Analysis code is available at https://github.com/fish-quant/autofish-analysis

## Results

Here, we present autoFISH (**Figure 1**), a universal toolbox to perform sequential smFISH with instrumentation that can be affordably built in house and adapted to specific needs and experimental protocols using standard buffers. autoFISH is based on several previous studies (Bintu et al., 2018; Imbert et al., 2022; Kishi et al., 2019; Moffitt and Zhuang, 2016; Tsanov et al., 2016) and is entirely open-source. We also provide analysis workflows in Python to analyze these data and infer single RNA quantification and the single cell level. autoFISH is hosted on GitHub (**see Methods**) with detailed building plans, control software, and experimental protocols. To validate and demonstrate the capability of autoFISH, we implemented and validated different protocols including workflows for signal amplification and tissue clearing on both cultured cells and tissue sections.

**Figure 1.**
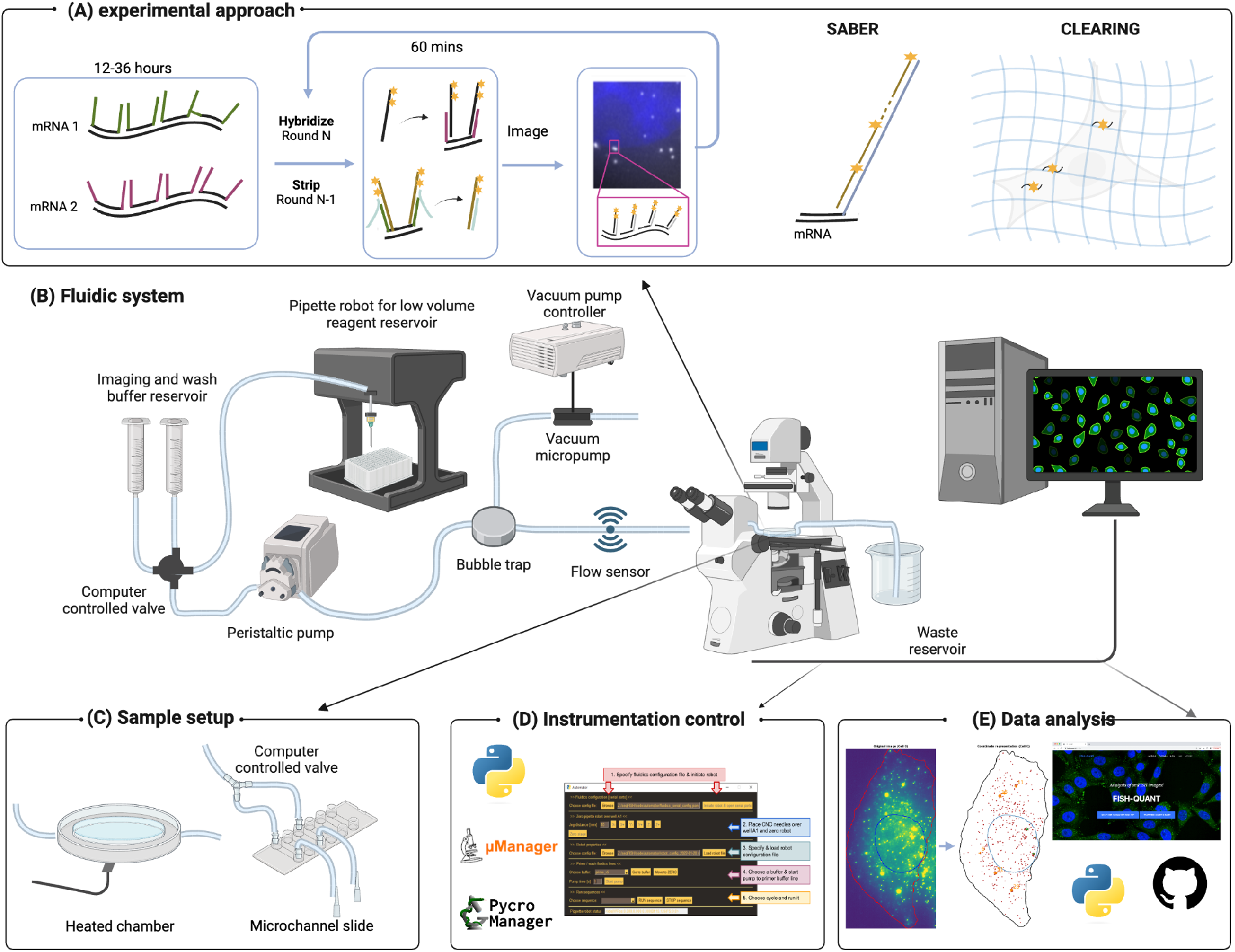
Overview of autoFISH summary. **A** Schematic of different experimental approaches. (Left) primary oligos against all target genes are hybridized simultaneously on the bench. In each of the N hybridization rounds, the secondary oligos revealing a subset of the RNA species is hybridized. Concurrently, stripping oligos against the previously imaged RNA can be hybridized. SABER provides moderate signal amplification by providing several binding sites for imager oligos. For clearing, RNAs are anchored into a polymer and the sample is then cleared to remove autofluorescence and scaffolds for non-specific oligo binding. **B** Overview of fluidics system with the different components. **C** Samples can either be placed in an imaging chamber (for tissues) or in disposable Ibidi microchannel slides (for cell). **D** Python control software allows the fluidics system and initiate communication with the microscope. **E** AProvided analysis software analysis different hybridization rounds and provide RNA abundance measurements for each segmented cell. Figure was created with BioRender.com.

### Computer controlled fluidics system for automated experiments

Sequential FISH experiment consists of several key steps: hybridization of primary oligos overnight at 37C, sample mounting onto the microscope followed by alternating imaging steps and incubation steps of fluorescently labeled secondary probe hybridization and displacement of secondary probes from the previous round (**Figure 1A**).

The central part of autoFISH is a computer-controlled fluidics system to perform the sequential hybridization steps and imaging automatically. This system allows flushing of buffers from syringes providing large buffer reservoirs or single-use buffers stored in multiwell plates accessed through a pipette robot. A computer controlled valve allows to select buffers and flow control is achieved through a peristaltic pump, with flow rates being monitored with a flow sensor. Lastly, air bubbles are removed from the buffers with a bubble trap. Samples can be mounted using an imaging chamber, ideal for single condition cell culture or tissue samples, or through an Ibidi microchannel slide, where up to six experimental conditions can be probed simultaneously (**Figure 1C**).

We want to emphasize two important aspects in our design. First, we found that placing the pump right after the valve yielded more robust performance than placing it after the imaging chamber (at the end of the fluidic system). Based on our experience, this resulted in a higher efficiency of the bubble trap as well as reduced deformation of the imaging chamber during pump action, which otherwise can result in loss of microscope autofocus. Second, we found that using Ibidi channel slides provided a more affordable alternative to the commonly used FCS2 imaging chamber, and enabled increasing the throughput of experiments when working with cultured cells, with only a moderate increase in buffer volumes. To guarantee homogenous equal flow through all channels, we added an outlet distribution valve to flow liquid through each channel consecutively. This leads to only marginal increases in the overall experimental timescale, but significantly improves the robustness of the system, especially compared to “static” one-to-many distribution where we found large variability in flow rates varied across channels, likely due to small differences in resistance of the fluidic paths.

We implemented a modular, open-access Python package that can control all components of the fluidic setup permitting to perform automated buffer exchanges. The modularity of the code makes it easy to add new components from other providers, e.g. to add a different peristaltic pump. The user can flexibly define fluidics runs in a human-readable text file containing all buffers, pump and incubation times, as well as optional steps, e.g. to perform a DAPI staining only for one run, or to add hybridization steps, e.g. for signal amplification with SABER.

For fully automated experiments, imaging has to be automated as well. As a design choice, we do not directly control microscopes from within our code, but provide different methods to interface with acquisition software packages. Currently, we support acquisition in micromanager via Pycromanger and triggered acquisition either with a TTL pulse or a shared synchronization text file (see **Methods** for details). With these solutions, autoFISH can already easily be used with different imaging systems, and new trigger approaches can easily be added if necessary (**Figure 1D**).

Lastly, we provide a graphical user interface (GUI) to facilitate usage. This interface can be used to control the entire workflow: load the configuration files, initiate the fluidics system, initiate trigger with the microscope, and then launch either an individual fluidics run or a fully automated run, where imaging and fluidics runs are performed sequentially and iteratively.

### autoFISH with 20 hybridization rounds

We first demonstrated the robustness of the system in generating specific smFISH signals and the efficiency of signal removal. We therefore performed an experiment consisting of 20 rounds where we alternated probing two genes whose RNA have distinct subcellular localization patterns: XPO1, which is randomly distributed, and KIF1, which preferentially localizes in cell protrusions (Chouaib et al., 2020). The same RNAs can be clearly seen in the respective first and last hybridization round for either gene (**Figure 2A**). Importantly, probe removal is efficient as the stripped signal from previous rounds is not visible (**Figure 2B**). More detailed analysis revealed a consistent number of detected spots per round (**Figure 2C**). Importantly, the detected spots were also the same, as indicated by a high colocalization percentage (approx 90%, **Figure 2D-E**, and **Supplementary Figure 1**), which is inline with other studies performing dual-color labeling of the same RNA (Tsanov et al., 2016). To evaluate signal conservation, we calculated the signal to noise ratio for each round. It remained constant (**Figure 2F**). Closer inspection revealed that the subsequent washing rounds slightly reduced the actual RNA signal, but also the background (**Figure 2G-H**). In summary, we demonstrated robust RNA detection over 20 hybridization rounds. By using multiple distinct fluorophores this allows to visualize several tens of RNAs.

**Figure 2.**
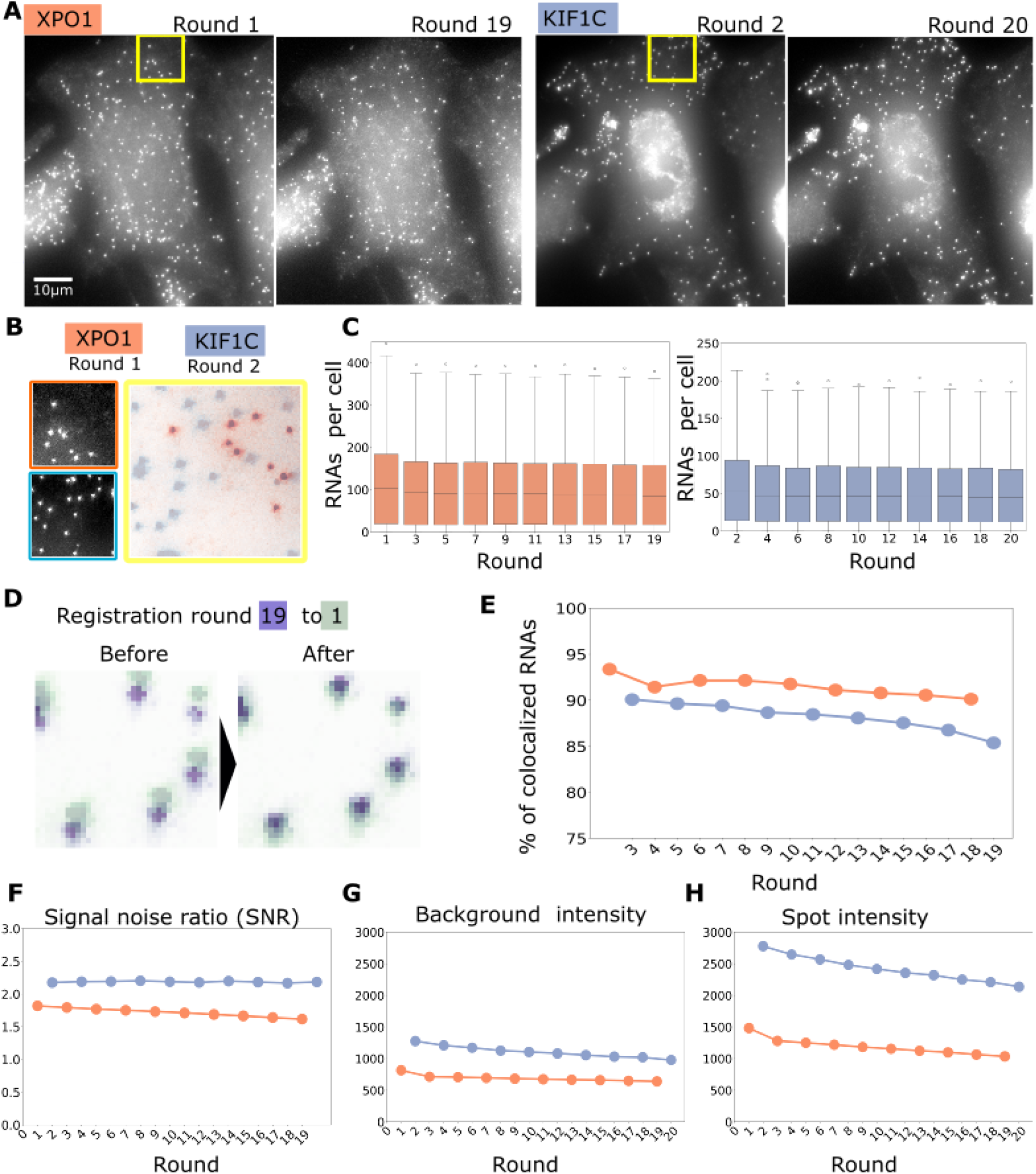
20 hybridization rounds with autoFISH. **A** Images of the same cell with either XPO1 (right) and KIF1c (left) for the respective first and last hybridization rounds. Same Intensity scaling [min, max] was used for both images of each gene: XPO1 [374,2216], KIF1c [2334,5708]. **B** Zooms in (yellow rectangles in A) showing XPO1 (round 1) and KIF1C (round 20). **C** RNA counts per cell across round for XPO1 (right) and KIF1C (left). **D** Example of registration with the plot of images from round 1 (green) and round 19 (red). **E** Pourcentage of colocalized RNAs with reference to the first respective hybridization round. **F-G-H** Mean Signal over noise ratio (SNR), background intensity and spots intensity across rounds.

### Flexible signal amplification with SABER

In smFISH, each RNA is targeted by several oligos. This leads to a strong local signal at each RNA that can be detected under a microscope. Signal amplification can help to increase RNA detection efficiency for more challenging samples. In autoFISH, we place readout sequences on both 3’and 5’ of the primary oligos. Further, each imaging oligos carries two fluorophores. So each primary oligos is labeled with up to 4 fluorophores. If the obtained signal is not strong enough, we occasionally also label the secondary oligos with two fluorophores. However, this is rather costly. An elegant alternative is SABER, which provides a robust and programmable solution (Kishi et al., 2019). In short, it allows the extension of oligos with a repeated sequence in-vitro before being used in hybridization (**Figure 3A**). These extended oligos can then be bound by several imaging oligos leading to an amplified signal. This extension can be precisely controlled by adjusting the duration of the amplification reaction (**Figure 3A**). A major advantage of performing the amplification before hybridization is that the duration of the actual FISH experiment is only minimally affected.

**Figure 3.**
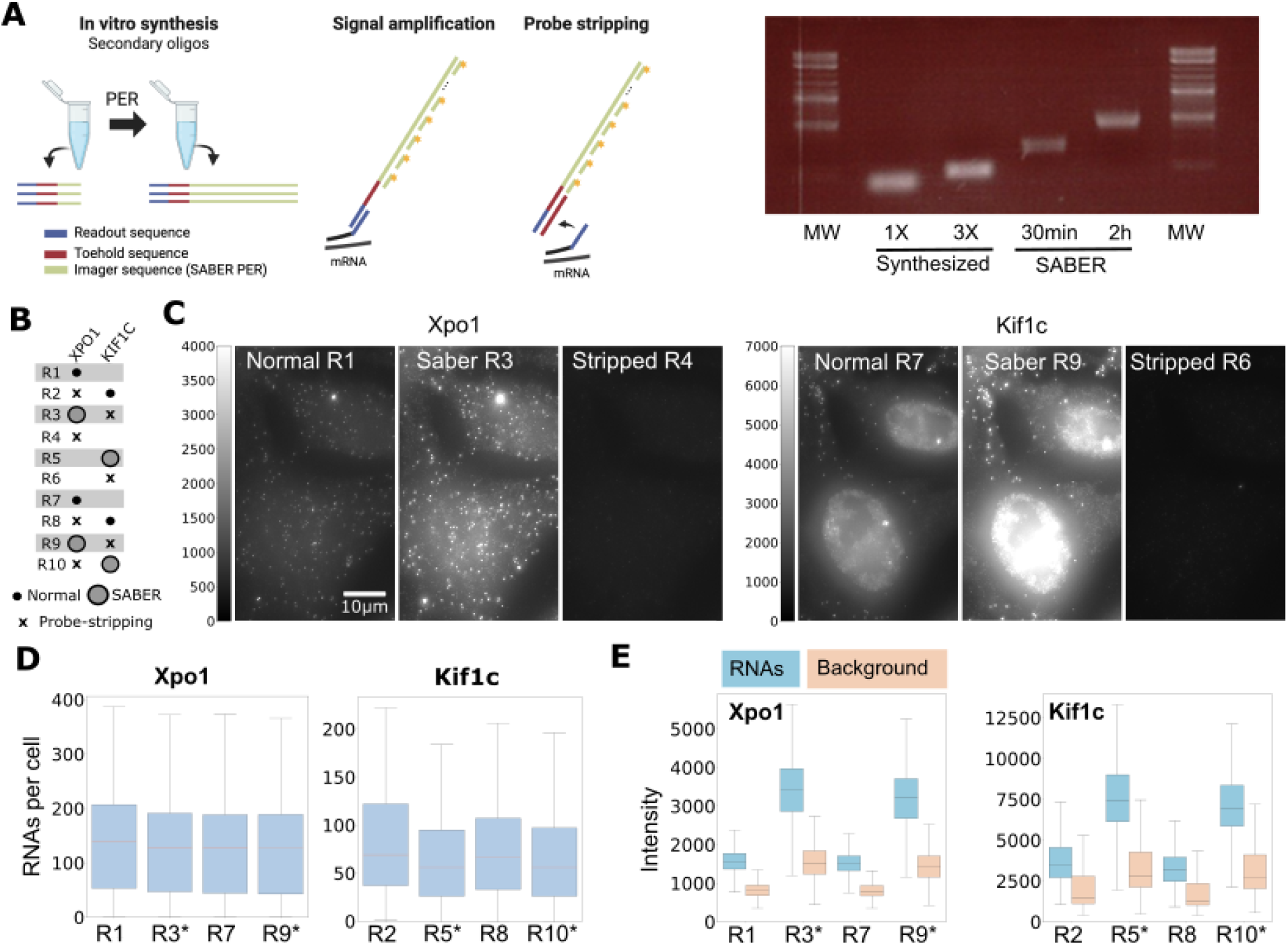
Signal amplification with SABER. **A** SABER allows to amplify a specific sequence acting as a binding region for imaging oligos. Hybridization of these oligos permits to amplify the signal. The oligo also contains a toehold region (red) which permits probe-stripping. Gel shows readout oligos with either 1 or 3 repeats or SABER amplification with two different durations (30 mins and 2h). **B** Hybridization conditions for the 10 rounds (R1-R10): small, filled dot indicates a standard FISH, larger dot indicates SABER FISH, x indicates stripping of the previous round. **C** Images of the same cell across different hybridization rounds. Images for the same gene are shown with the same intensity scaling. **D** RNA levels per cell across rounds. SABER runs are indicated with an asterisk. **E** Quantification of intensities of RNA and background across rounds. SABER runs are indicated with an asterisk.

We integrated SABER in autoFISH by replacing the imager sequence on the readouts by the PER sequence of SABER required for amplification (**Figure 3A**). We then optimized the protocol to be compatible with the fluidics system. In short, we replaced the formamide from the original buffers with the less toxic Ethylene Carbonate. We also performed the hybridization of the SABER oligos in two steps on the fluidics system: first the amplicons and then the imaging oligos. We performed the hybridization of the amplicons in a more stringent buffer which leads to substantial decrease in background. We also found that adding even a moderate amount of dextran (5% was compatible with our fluidics system) reduced the required amount of imaging oligo therefore reducing experimental cost. With these changes, SABER can be performed on the fluidics system. Interestingly, the flexibility in the fluidics control allows us to perform SABER hybridization, which requires an additional hybridization step, in selected runs only as demonstrated in the validation experiments.

We then performed an experiment targeting XPO1 and KIF1C and alternated between normal and SABER hybridization, and also included a control round where we only stripped SABER without a new hybridization (**Figure 3B**). Qualitative inspection of the images shows a clear amplification of SABER for either gene with the same RNAs being visible and SABER leading to an increased signal intensity (**Figure 3B**). Importantly, the SABER amplicon can also be efficiently stripped and the signal removed (**Figure 3B)**. More detailed analysis confirmed this qualitative impression, with a constant amount of detected RNA per round (**Figure 3C**). This analysis also revealed that SABER increased RNA intensity **Figure 3D**, but also a slight increase in background (**Figure 3D)**.

These experiments demonstrate how SABER can be used in autoFISH for signal amplification. Further, it showcasts the flexibility in the control software with different hybridization conditions for specific runs.

### autoFISH on cleared mouse lung tissue

In a recent study, we performed smFISH of selected RNA markers on mouse lung tissue (Curras-Alonso et al., 2023). While imaging quality was sufficient for analysis, we also encountered substantial background fluorescence. It has been shown that this background stems both from tissue autofluorescence and non-specific binding of probes to different cellular structures (Moffitt et al., 2016a). Tissue clearing methods help to efficiently reduce this background. Here, the biological sample is embedded into a nonswellable polyacrylamide (PA) matrix (**Figure 4A**). Several strategies to anchor RNAs to the PA exist, either by chemical compounds (Chen et al., 2016) or the more widely used approach by using oligos targeting the Poly-A tails of mRNAs carrying a terminal acrydite moeity (Moffitt et al., 2016a). The tissue can then be treated with ProteinaseK and SDS to remove proteins and lipids which are the major cause for autofluorescence and non-specific binding (Moffitt et al., 2016a).

**Figure 4.**
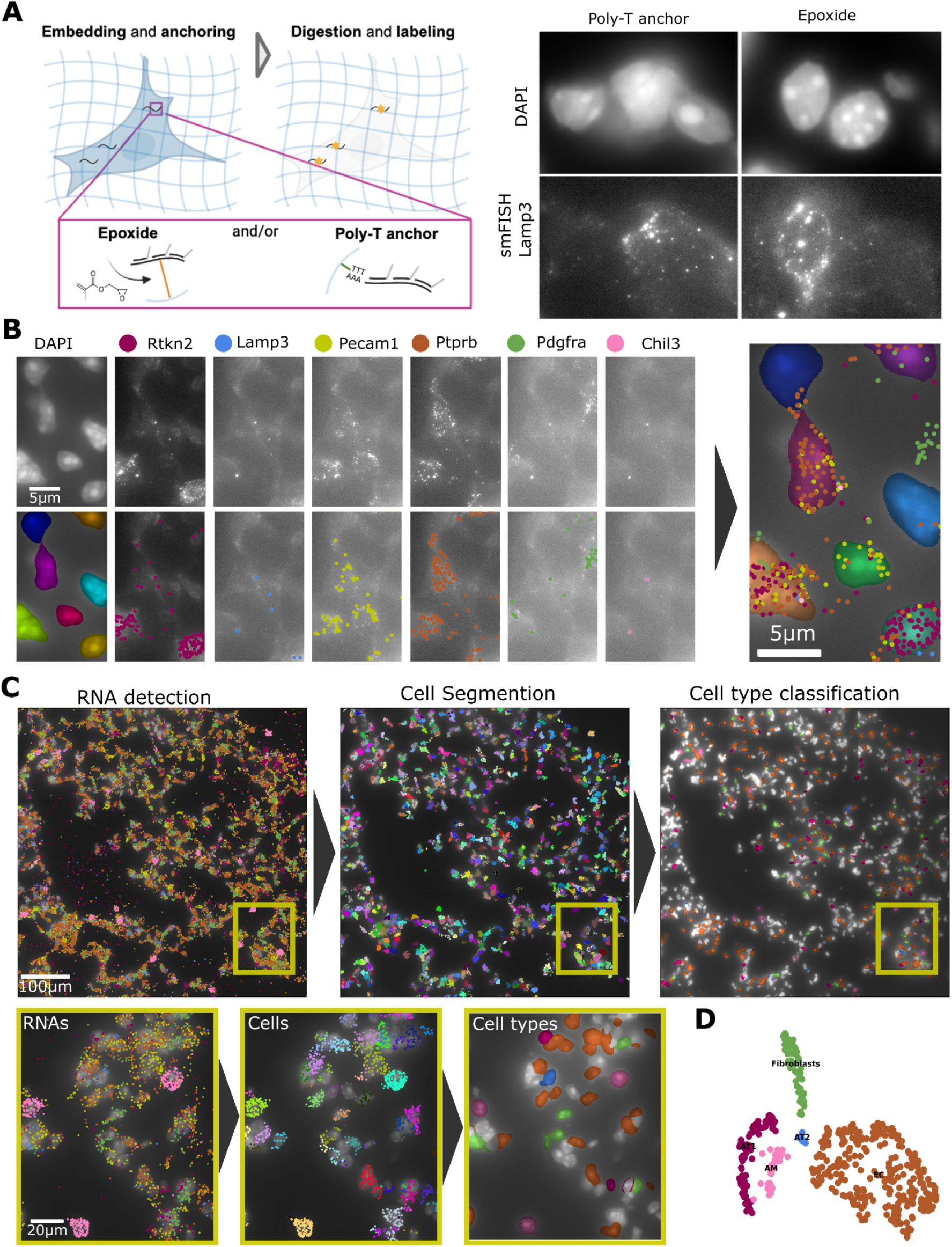
autoFISH on cleared mouse lung tissue. **A** (Left) Principle of clearing. RNAs can be anchored with epoxide and/or Poly-T oligos carrying an acrydite moeity. Panel was created with BioRender.com. (Right) Influence of epoxide on nuclear signals. DAPI and smFISH images after clearing with Poly-T anchoring (left) and epoxide (right). **B** Nucleus segmentation and RNA spots detection across rounds. **C** Illustration of workflow from RNA detection, to cell segmentation and cell type classification. Yellow rectangle shows a region provided with a zoom-in. **D** Cell-type mapping based on our reference scRNAseq data.

When applying the protocol with the Poly-T anchoring oligos to lung tissue, background was reduced and RNAs were detected, but we found that the nuclear signal was often fuzzy compared to samples anchored with epoxide (**Figure 4A**). For samples such as lung tissue with locally high density of nuclei, this could then have a decremental impact on nuclear segmentation. We speculated that this could be due to the reduced constraints of chromatin after digestion. In a recent study, an epoxide (GMA) was introduced as a universal linking agent in Expansion microscopy (ExM) for proteins and RNA (Cui et al., 2023). Since polymers used for clearing and ExM are similar, we speculated that such an epoxide could be used for clearing protocols as well. When used alone or combined with the poly-T anchor oligos, we found indeed that nuclear staining appears more confined (**Figure 4A**). We then performed autoFISH against 6 targets from our previous study, where we detect individual RNAs of each RNA species (**Figure 4B**). Since we did not use dedicated membrane-markers, we applied our analysis package ComSeg (Defard et al., 2024b) to perform cell segmentation leveraging nuclei staining and the detected RNA point clouds (**Figure 4C**). The resulting gene expression profiles then allow clustering (**Figure 4D**) and mapping the identified cell-types back on the image, revealing their spatial location.

In summary, we introduced an updated tissue clearing procedure, where the usage of epoxide leads to better retention of the nuclear signal. We successfully applied this approach to mouse tissue to detect different marker genes.

## Discussion

smFISH is a powerful technique for visualizing individual RNAs within their native cellular context. By performing multiple cycles of probe hybridization, imaging, and probe removal, the number of RNA species that can be analyzed is greatly increased. However, this also requires more complex protocols and specialized equipment. Although commercial solutions are available, they are costly and lack flexibility. Here, we introduce autoFISH, a modular toolkit designed for automated, sequential FISH, utilizing validated and optimized workflows.

In this work, we demonstrate the performance of autoFISH both in cell lines and tissue with up to 20 hybridization runs. While the throughput of this method is lower than that of commercial multiplexed solutions, it offers some advantages. Firstly, since each RNA is imaged individually, spatial crowding is not a concern, allowing even highly expressed genes to be targeted. Additionally, by ordering oligos for each gene separately, we can customize the selection of targeted genes. One of the main strengths of autoFISH is its flexibility. We demonstrate this by applying various experimental protocols, showcasing flexible control over fluidics runs through optional steps, and implementing different methods to trigger acquisitions on the microscope.

For more challenging samples, we provide experimental protocols for signal amplification and tissue clearing that are compatible with the autoFISH fluidic system. We illustrate how signal amplification can be achieved with SABER, and show that such amplification can be performed on specific runs due to the flexibility in designing fluidic runs. The proposed use of epoxide in tissue clearing enhances nucleus signal preservation, promising to improve segmentation in densely packed samples. Additionally, the epoxide used (GMA) is inexpensive, allowing for cost-effective experiments. Finally, epoxide anchoring is more versatile than targeting poly-A tails, which limits the application to mature mRNAs only. This versatility ensures that other RNA species, such as nascent or long non-coding RNAs, can also be anchored and are less likely to be washed out during the clearing step.

We offer several options to trigger acquisitions, with the potential to add more in the future, making it straightforward to integrate autoFISH in different imaging systems. We also demonstrate the use of different sample supports, allowing for imaging of tissue sections or cell lines in different conditions. The open design of the control software facilitates the addition of new components, such as valves or pumps. The control software is implemented in Python, enabling easy implementation of additional steps, such as quality control, and leveraging powerful libraries within the Python ecosystem.

In summary, we believe that autoFISH will lower the access barrier for research groups to perform sequential FISH. We believe that in-house solutions, such as the one presented here, are also well suited to be used side-by-side with commercial approaches. The latter permit to perform large-scale high-multiplexed experiments that are very challenging. Appropriate analysis of these and other data (such as scRNAseq) can then reveal a smaller number of target genes. With in-house solutions these genes can be probed at substantially reduced cost. Lastly, autoFISH is developed as Open Source, and we hope that it will spark community effort and will be expanded with more protocols and modules from us and other contributors.

## Acknowledgements

This work has received financial support through the Institut Pasteur (Single Cell Seed Grant), Agence Nationale de la Recherche (ANR) grants LUSTRA (ANR-19-CE14-0015-03) and TRANSFACT (ANR-19-CE12-0007-02). T.D, F.M. and C.W. acknowledge funding by Institut Pasteur. A.M. has received PhD fellowship from the European Union’s Horizon 2020 research and innovation program under the Marie Skłodowska-Curie grant agreement No 847718. H.L. is the recipient of a PhD fellowship from the International Student program from Paris-Saclay University.

## References

Beare, R., Lowekamp, B., Yaniv, Z., 2018. Image Segmentation, Registration and Characterization in R with SimpleITK. J Stat Softw 86, 8. 10.18637/jss.v086.i08

Bintu, B., Mateo, L.J., Su, J.-H., Sinnott-Armstrong, N.A., Parker, M., Kinrot, S., Yamaya, K., Boettiger, A.N., Zhuang, X., 2018. Super-resolution chromatin tracing reveals domains and cooperative interactions in single cells. Science 362. 10.1126/science.aau1783

Chen, F., Wassie, A.T., Cote, A.J., Sinha, A., Alon, S., Asano, S., Daugharthy, E.R., Chang, J.-B., Marblestone, A., Church, G.M., Raj, A., Boyden, E.S., 2016. Nanoscale imaging of RNA with expansion microscopy. Nat. Methods 13, 679–684. 10.1038/nmeth.3899

Chouaib, R., Safieddine, A., Pichon, X., Imbert, A., Kwon, O.S., Samacoits, A., Traboulsi, A.-M., Robert, M.-C., Tsanov, N., Coleno, E., Poser, I., Zimmer, C., Hyman, A., Le Hir, H., Zibara, K., Peter, M., Mueller, F., Walter, T., Bertrand, E., 2020. A Dual Protein-mRNA Localization Screen Reveals Compartmentalized Translation and Widespread Co-translational RNA Targeting. Dev Cell 54, 773-791.e5. 10.1016/j.devcel.2020.07.010

Cui, Y., Yang, G., Goodwin, D.R., O’Flanagan, C.H., Sinha, A., Zhang, C., Kitko, K.E., Shin, T.W., Park, D., Aparicio, S., Boyden, E.S., 2023. Expansion microscopy using a single anchor molecule for high-yield multiplexed imaging of proteins and RNAs. PLoS One 18, e0291506. 10.1371/journal.pone.0291506

Curras-Alonso, S., Soulier, J., Defard, T., Weber, C., Heinrich, S., Laporte, H., Leboucher, S., Lameiras, S., Dutreix, M., Favaudon, V., Massip, F., Walter, T., Mueller, F., Londoño-Vallejo, J.-A., Fouillade, C., 2023. An interactive murine single-cell atlas of the lung responses to radiation injury. Nat Commun 14, 2445. 10.1038/s41467-023-38134-z

Defard, T., Desrentes, A., Fouillade, C., Mueller, F., 2024a. A DIY guide for image-based spatial transcriptomic: TLS as a case example. bioRxiv 2024–07.

Defard, T., Laporte, H., Ayan, M., Soulier, J., Curras-Alonso, S., Weber, C., Massip, F., Londoño-Vallejo, J.-A., Fouillade, C., Mueller, F., Walter, T., 2024b. A point cloud segmentation framework for image-based spatial transcriptomics. Commun Biol 7, 823. 10.1038/s42003-024-06480-3

Eng, C.-H.L., Lawson, M., Zhu, Q., Dries, R., Koulena, N., Takei, Y., Yun, J., Cronin, C., Karp, C., Yuan, G.-C., Cai, L., 2019. Transcriptome-scale super-resolved imaging in tissues by RNA seqFISH. Nature 568, 235–239. 10.1038/s41586-019-1049-y

Imbert, A., Ouyang, W., Safieddine, A., Coleno, E., Zimmer, C., Bertrand, E., Walter, T., Mueller, F., 2022. FISH-quant v2: a scalable and modular tool for smFISH image analysis. RNA 28, 786–795. 10.1261/rna.079073.121

Kishi, J.Y., Lapan, S.W., Beliveau, B.J., West, E.R., Zhu, A., Sasaki, H.M., Saka, S.K., Wang, Y., Cepko, C.L., Yin, P., 2019. SABER amplifies FISH: enhanced multiplexed imaging of RNA and DNA in cells and tissues. Nat Methods 16, 533–544. 10.1038/s41592-019-0404-0

Lohoff, T., Ghazanfar, S., Missarova, A., Koulena, N., Pierson, N., Griffiths, J.A., Bardot, E.S., Eng, C.-H.L., Tyser, R.C.V., Argelaguet, R., Guibentif, C., Srinivas, S., Briscoe, J., Simons, B.D., Hadjantonakis, A.-K., Göttgens, B., Reik, W., Nichols, J., Cai, L., Marioni, J.C., 2022. Integration of spatial and single-cell transcriptomic data elucidates mouse organogenesis. Nat Biotechnol 40, 74–85. 10.1038/s41587-021-01006-2

Mateo, L.J., Sinnott-Armstrong, N., Boettiger, A.N., 2021. Tracing DNA paths and RNA profiles in cultured cells and tissues with ORCA. Nat Protoc 16, 1647–1713. 10.1038/s41596-020-00478-x

Moffitt, J.R., Hao, J., Bambah-Mukku, D., Lu, T., Dulac, C., Zhuang, X., 2016a. High-performance multiplexed fluorescence in situ hybridization in culture and tissue with matrix imprinting and clearing. Proc. Natl. Acad. Sci. U.S.A. 10.1073/pnas.1617699113

Moffitt, J.R., Hao, J., Wang, G., Chen, K.H., Babcock, H.P., Zhuang, X., 2016b. High-throughput single-cell gene-expression profiling with multiplexed error-robust fluorescence in situ hybridization. Proc. Natl. Acad. Sci. U.S.A. 113, 11046–11051. 10.1073/pnas.1612826113

Moffitt, J.R., Zhuang, X., 2018. RNA Imaging with MERFISH - Probe Construction.

Moffitt, J.R., Zhuang, X., 2016. RNA Imaging with Multiplexed Error-Robust Fluorescence In Situ Hybridization (MERFISH). Meth. Enzymol. 572, 1–49. 10.1016/bs.mie.2016.03.020

Pachitariu, M., Stringer, C., 2022. Cellpose 2.0: how to train your own model. Nat Methods 1–8. 10.1038/s41592-022-01663-4

Palla, G., Fischer, D.S., Regev, A., Theis, F.J., 2022. Spatial components of molecular tissue biology. Nat Biotechnol 40, 308–318. 10.1038/s41587-021-01182-1

Pichon, X., Lagha, M., Mueller, F., Bertrand, E., 2018. A Growing Toolbox to Image Gene Expression in Single Cells: Sensitive Approaches for Demanding Challenges. Mol. Cell 71, 468–480. 10.1016/j.molcel.2018.07.022

Pinkard, H., Stuurman, N., Ivanov, I.E., Anthony, N.M., Ouyang, W., Li, B., Yang, B., Tsuchida, M.A., Chhun, B., Zhang, G., Mei, R., Anderson, M., Shepherd, D.P., Hunt-Isaak, I., Dunn, R.L., Jahr, W., Kato, S., Royer, L.A., Thiagarajah, J.R., Eliceiri, K.W., Lundberg, E., Mehta, S.B., Waller, L., 2021. Pycro-Manager: open-source software for customized and reproducible microscope control. Nat Methods 18, 226–228. 10.1038/s41592-021-01087-6

Preibisch, S., Saalfeld, S., Tomancak, P., 2009. Globally optimal stitching of tiled 3D microscopic image acquisitions. Bioinformatics 25, 1463–1465. 10.1093/bioinformatics/btp184

Rademacher, A., Huseynov, A., Bortolomeazzi, M., Wille, S.J., Schumacher, S., Sant, P., Keitel, D., Okonechnikov, K., Ghasemi, D.R., Pajtler, K.W., Mallm, J.-P., Rippe, K., 2024. Comparison of spatial transcriptomics technologies using tumor cryosections. 10.1101/2024.04.03.586404

Tsanov, N., Samacoits, A., Chouaib, R., Traboulsi, A.-M., Gostan, T., Weber, C., Zimmer, C., Zibara, K., Walter, T., Peter, M., Bertrand, E., Mueller, F., 2016. smiFISH and FISH-quant - a flexible single RNA detection approach with super-resolution capability. Nucleic Acids Res. 44, e165. 10.1093/nar/gkw784

Xu, W., Cai, H., Zhang, Q., Mueller, F., Ouyang, W., Cao, G., 2024. U-FISH: a universal deep learning approach for accurate FISH spot detection across diverse datasets. 10.1101/2024.03.06.583706

Zhang, M., Eichhorn, S.W., Zingg, B., Yao, Z., Cotter, K., Zeng, H., Dong, H., Zhuang, X., 2021. Spatially resolved cell atlas of the mouse primary motor cortex by MERFISH. Nature 598, 137–143. 10.1038/s41586-021-03705-x

